# Peripheral sensory terminal degeneration is linked to sensory fiber hyperexcitability in paclitaxel-induced peripheral neuropathy

**DOI:** 10.64898/2026.07.15.738514

**Authors:** Junhui Du, Abdi Salahadin, Zizhen Wu, Qing Yang

## Abstract

Paclitaxel-induced peripheral neuropathy (PIPN) is the most common dose-limiting side effect of paclitaxel chemotherapy, yet the relationship between structural damage to peripheral nerve terminals and functional impairment of nociceptors has not been directly examined. Here we combined *ex vivo* skin-nerve electrophysiology with three-dimensional (3D) imaging of fDISCO-cleared glabrous skin to characterize both morphological and functional changes in peripheral sensory terminals following paclitaxel treatment in rats. Five weeks after paclitaxel administration, skin-nerve recordings revealed a marked increase in the proportion of C- and Aδ-fibers exhibiting spontaneous discharge (approximately 2.4- and 2.6-fold increases, respectively) compared to vehicle-treated controls. Paclitaxel also selectively reduced the mechanical activation threshold of C-fibers without affecting Aδ fibers. Three-dimensional reconstruction of PGP9.5-immunolabeled nerve terminals showed that sensory fibers in glabrous skin form a vertically oriented, tree-like architecture. Paclitaxel treatment severely reduced both the terminal branch length and the density of free nerve endings in the epidermis. Strikingly, co-administration of the Kv7 channel activator retigabine prevented both the electrophysiological and morphological alterations induced by paclitaxel. These findings provide direct evidence that peripheral nerve terminal degeneration and hyperexcitability co-occur in PIPN and that Kv7 channel activation can protect against both structural and functional damage.

**Significance:** This study provides the first direct evidence linking intraepidermal sensory terminal degeneration with peripheral sensory fiber hyperexcitability in paclitaxel-induced peripheral neuropathy by combining skin–nerve electrophysiology with three-dimensional imaging of sensory terminals. Paclitaxel induces distal degeneration of intraepidermal sensory terminals, increases spontaneous discharge, and lowers the mechanical activation threshold of C-fibers, demonstrating that structural degeneration and functional abnormalities occur concurrently in the peripheral terminal. Retigabine prevents both structural and functional alterations, supporting peripheral sensory terminals as a therapeutic target for preventing chemotherapy-induced neuropathy.

## INTRODUCTION

Paclitaxel-based chemotherapy is widely used to treat breast and ovarian cancers, but peripheral neurotoxicity remains its major dose-limiting side effect. Paclitaxel-induced peripheral neuropathy (PIPN) presents as a predominant sensory dysfunction, often accompanied by numbness and neuropathic pain, affecting fibers in a characteristic “stocking-and-glove” distribution ^10, 16^. Symptoms develop within days to months of treatment initiation and may persist long after chemotherapy ends ^12^. The incidence is strongly dose-dependent, with up to 84% of patients experiencing some degree of neuropathy ^12^. Despite progress in understanding the molecular mechanisms of PIPN, the relationship between structural nerve damage and functional impairment remains poorly characterized.

The hallmark of PIPN is the degeneration of intraepidermal nerve fibers (IENFs) ^5, 15, 33^. Both morphological and electrophysiological approaches have been used to study how IENF loss contributes to PIPN symptoms ^5, 15, 33, 34, 36^. Electrophysiological recordings from dorsal root ganglion (DRG) somata and sural nerves show increased ectopic spontaneous activity after paclitaxel treatment, yet these recording sites lack corresponding morphological changes ^34, 36^. Multiple sites along primary sensory neurons can generate ectopic activity after damage ^4, 17^, but whether the peripheral nerve terminals themselves are the origin of altered excitability after paclitaxel remains unknown. Furthermore, both C- and Aδ-fiber free endings are present at the dermal-epidermal junction ^18^, yet conventional IENF staining only reveals vertically oriented fibers in 2D histological sections. How these nerve terminals are distributed three-dimensionally, and how their spatial organization changes after chemotherapy, has not been examined. This gap is particularly important because mechanical sensitivity is often heightened even as intraepidermal free nerve ending density declines.

In this study, we combined *ex vivo* skin-nerve electrophysiology with three-dimensional morphological analysis to re-examine the relationship between IENF integrity and nociceptor function following paclitaxel treatment. Skin-nerve recordings allow mechanical stimulation to be applied directly to the skin, overcoming technical limitations of previous approaches ^34, 36^. IENFs were visualized in three dimensions using fDISCO tissue clearing, providing a comprehensive view of peripheral nerve terminal architecture. Because our previous work showed that paclitaxel initiates PIPN by inhibiting Kv7 ion channels in primary sensory neurons ^33^, we also tested whether co-administration of the Kv7 channel activator retigabine could prevent the morphological and electrophysiological changes induced by paclitaxel.

We found that paclitaxel markedly increased nociceptor excitability in skin-nerve preparations. Three-dimensional imaging revealed that sensory fibers in glabrous skin form a tree-like architecture, and that vertically oriented terminal branches were severely reduced after paclitaxel, consistent with decreased IENF density. Co-administration of retigabine attenuated both the electrophysiological and morphological changes. These results provide direct evidence linking terminal morphological damage to functional alterations in PIPN and support Kv7 channels as a therapeutic target.

## MATERIALS AND METHODS

All experimental procedures were conducted in accordance with the guidelines established by the International Association for the Study of Pain (IASP) and were approved by the Institutional Animal Care and Use Committee (IACUC) of the University of Texas Medical Branch at Galveston (UTMB). This study adheres to the ARRIVE 2.0 guidelines to ensure transparent and reproducible reporting. Adult male Sprague-Dawley rats (250-400 g) from Charles River Laboratory were used throughout this study. A total of 49 animals were included across all experimental groups. Animals were group-housed in pairs on wood chip bedding in a temperature-controlled facility (21 ± 1 °C) under a 12-hour light/dark cycle (lights on at 07:00). Standard laboratory chow and water were available ad libitum. Cages were cleaned and bedding replaced twice weekly. Upon arrival, all animals were allowed a one-week acclimation period before any experimental procedures.

### Drug administration

Paclitaxel (clinically formulated; TEVA Pharmaceuticals Inc., North Wales, PA) was dissolved in a vehicle stock solution consisting of an equal-volume mixture of Cremophor EL (Sigma-Aldrich, St. Louis, MO) and absolute ethanol, matching the clinical formulation. The stock solution was further diluted in sterile 0.9% saline immediately before injection. Paclitaxel was administered intraperitoneally (i.p.) at a dose of 2 mg/kg on Days 1, 3, 5, and 7 (four injections total, every other day), with all injections performed between 08:00 and 10:00 to minimize circadian variability. Injection volumes were adjusted according to individual body weight (e.g., a 250 g rat received 0.5 mL). Sham control animals received an equivalent volume of vehicle (Cremophor EL/ethanol diluted in 0.9% saline) on the same schedule.

Retigabine (Toronto Research Chemicals, North York, Ontario, Canada) was dissolved in sterile 0.9% saline and administered i.p. at 10 mg/kg twice daily from Day 1 through Day 10 ^15, 33^. On paclitaxel injection days (Days 1, 3, 5, and 7), the morning dose of retigabine was administered 30 minutes prior to paclitaxel injection to ensure adequate drug exposure. The second daily dose was given in the late afternoon (approximately 16:00-18:00). Animals in the paclitaxel-alone group received equivalent volumes of sterile 0.9% saline on the same twice-daily schedule to control for injection stress.

Animals were randomly assigned to one of three experimental groups prior to treatment onset: (1) Sham group (14 rats), receiving vehicle for paclitaxel and vehicle for retigabine on matching schedules; (2) Paclitaxel group (25 rats), receiving paclitaxel (2 mg/kg, i.p., Days 1, 3, 5, and 7) and vehicle of retigabine (i.p., twice daily, Days 1-10); (3) Paclitaxel + retigabine group (14 rats), receiving paclitaxel as above plus retigabine (10 mg/kg, i.p., twice daily, Days 1-10). Group assignment was performed using a randomization scheme to minimize selection bias, and experimenters were blinded to group allocation during behavioral assessments and electrophysiological recordings.

### Skin-nerve preparation

The *ex vivo* glabrous skin-nerve preparation was adapted from the protocol originally established in Carlton’s laboratory ^13^, with minor modifications. Five weeks after the final paclitaxel injection, glabrous skin was dissected from the plantar surface of the hindpaw, extending from the ankle to the distal tips of the toes, with a narrow border of adjacent hairy skin retained to prevent edge-related tissue damage and maintain preparation integrity. The tibial nerve and its medial and lateral plantar branches (approximately 3 cm in total length) were carefully dissected free from surrounding connective tissue at the level of the ankle joint while preserving their continuity with the skin flap (**Fig. 1A**).

**Figure 1.**
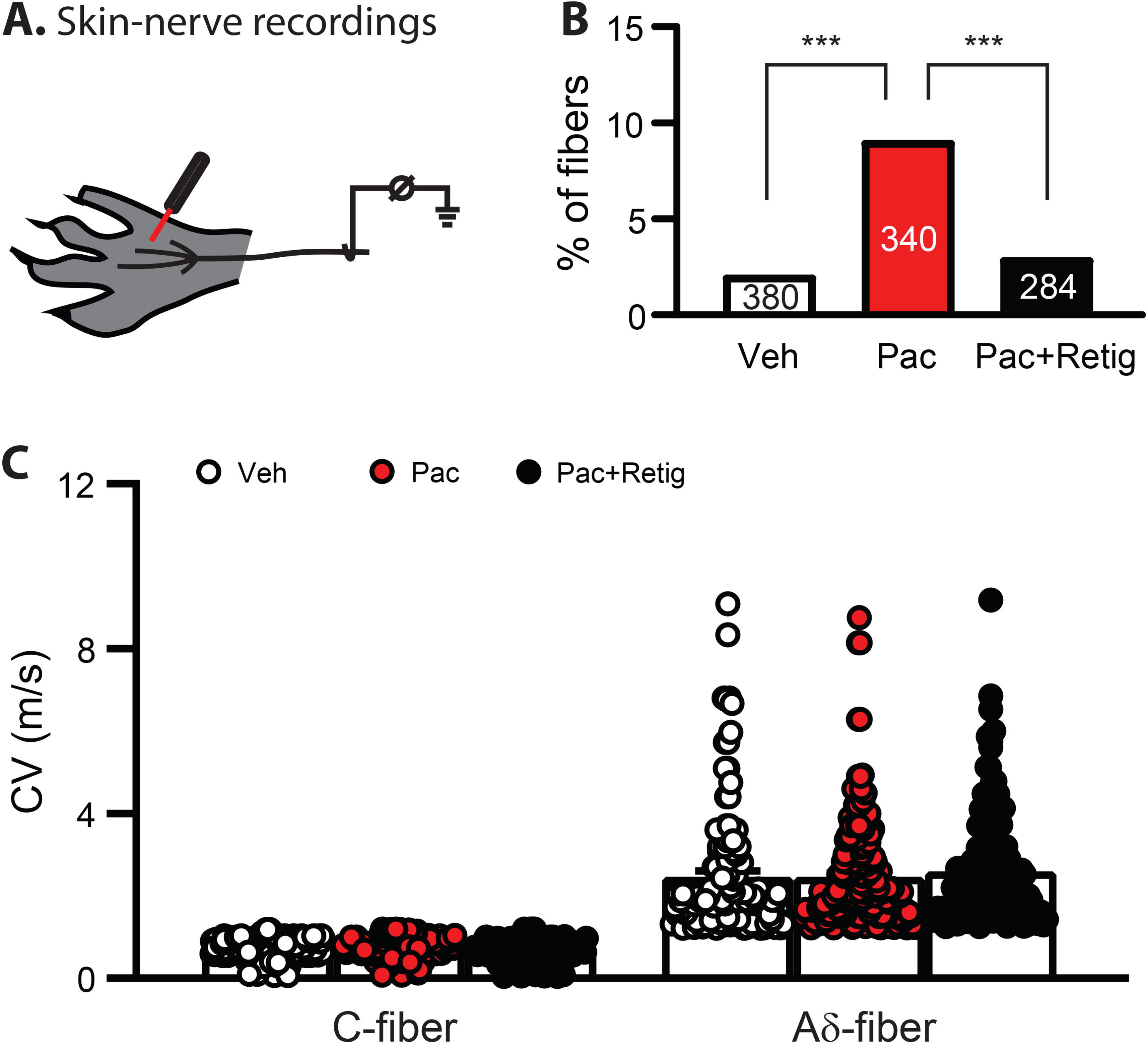
The effect of Paclitaxel treatment on conduction velocity (CV) of nociceptors. (**A**) Schematic drawing of the rat skin-nerve recording. Mechanical stimulation were applied sequentially to the skin with 5-minute rest periods in between each stimulus. (**B**) Percentage of fibers that are not responsive to electrical stimuli. Chi-square test, X^2^ = 22.09, P < 0.0001. The number in each column indicates the fibers recorded. (**C**) Scatter plots of the CV of all nociceptors recorded in Vehicle- (Veh), paclitaxel-treated (Pac), and paclitaxel plus retigabine (retig)-treated rats (columns represent mean ± SEM). Repeated measures one-way ANOVA followed by Tukey’s multiple comparison test (C-fibers, F_(2, 500)_ = 7.08, P = 0.0009; Veh vs Pac, P = 0.0734; Veh vs Pac+Retig, P = 0.0832. Aδ-fibers, F_(2, 450)_ = 0.45, P = 0.63378; Veh vs Pac, P = 0.9991, Veh vs Pac+Retig, P = 0.73). Each dot in column represents one fiber. Veh, vehicle; Pac, paclitaxel; Retig, retigabine.

The completed skin-nerve preparation was transferred to a custom organ bath and positioned corium-side up on a stainless-steel grid. The preparation was continuously superfused at 15 mL/min with modified synthetic interstitial fluid (SIF) maintained at 32 ± 0.5 °C and equilibrated with 95% O_2_/5% CO_2_. The ionic composition of the SIF was (in mM): NaCl 123, KCl 3.5, MgSO4 0.7, CaCl_2_ 2.0, sodium gluconate 9.5, NaH_2_PO_4_ 1.7, glucose 5.5, sucrose 7.5, and HEPES 10; pH was adjusted to approximately 7.4 with NaOH (osmolarity approximately 305 mOsm/kg).

The plantar nerve branches were guided through a small partition into an adjacent recording chamber, where they were submerged beneath a thin layer of mineral oil overlaying a base layer of SIF to maintain tissue hydration while providing an electrically isolated recording environment. On a mirrored dissection stage, nerve trunks were carefully desheathed under a stereodissecting microscope using sharpened watchmaker forceps (Dumont #5). Individual nerve filaments from the medial or lateral plantar branch were progressively teased into finer bundles until the smallest viable functional unit was isolated without mechanical rupture or loss of electrical activity. Filament viability was confirmed prior to each recording by verifying the presence of a detectable compound action potential in response to brief mechanical or electrical stimulation applied to the skin.

### Neurophysiological recording

Extracellular single-unit recordings were obtained from teased nerve filaments using gold wire electrodes connected to a differential amplifier (DAM80; World Precision Instruments, Sarasota, FL). Recorded signals were bandpass filtered (300 Hz - 3 kHz) and amplified prior to digitization. Action potentials were acquired and stored in real time via a CED1401 data acquisition interface (Cambridge Electronic Design Ltd., Cambridge, UK) and analyzed offline using Spike2 software (Cambridge Electronic Design) for spike sorting and waveform discrimination. Template-based waveform matching confirmed single-unit isolation throughout each recording session; units with waveform instability or signal-to-noise ratios below 3:1 were excluded.

### Fiber identification and classification

Following filament placement on the recording electrode, a waiting period was observed before recordings began to exclude possible “injury discharge,” which lasts no more than one to two minutes in the vast majority of fibers ^6, 20, 31^. A two-minute baseline of spontaneous activity was then recorded prior to any mechanical stimulation to assess background firing rate. Spontaneous discharge was defined as any activity exceeding 1 impulse/min ^3, 32^. Mechanically sensitive fibers were identified by applying focal mechanical stimulation across the skin surface using a blunt glass probe (approximately 1 mm tip diameter) to locate receptive fields. Only units exhibiting a clear and reproducible response to noxious stimulation (von Frey threshold not less than 5 mN) were selected for further characterization, excluding the possible inclusion of low-threshold slowly adapting mechanoreceptors.

Electrical stimulation of the receptive field was used to calculate conduction velocity (CV). CV was determined by monopolar electrical stimulation delivered at the most mechanosensitive point within the receptive field using a Teflon-coated stainless-steel electrode (impedance ∼2 MΩ), gently lowered into the skin at the receptive field center. Electrical pulses were delivered at 1 Hz with pulse durations of 0.3 to 3.0 ms and amplitudes of 0.02 to 1.0 mA, adjusted to reliably evoke a single action potential at minimal stimulus intensity. CV was calculated by dividing the conduction distance by the latency of the electrically evoked action potential measured from stimulus onset to the peak of the first deflection. Fibers were classified based on CV: Aδ-fibers had CVs between 1.0 and 10 m/s, and C-fibers had CVs between 0.2 and 1.0 m/s ^13^.

### Mechanical stimulation protocol

Following fiber identification and CV determination, the mechanical response properties of each unit were characterized using a controlled ramp-and-hold stimulus delivered via a feedback-controlled mechanical stimulator (Aurora Scientific, Aurora, Ontario, Canada). A motor-driven stylus (0.7 mm diameter) delivered mechanical forces perpendicular to the corium side in the most sensitive region of the receptive field (**Fig. 1A**). Mechanical threshold was defined as the minimum force (in mN) required to evoke at least two action potentials during the ascending ramp phase (0-170 mN over 20 s). Suprathreshold responses were assessed by delivering a square wave pulse (180 mN, 10 s), and firing frequency was quantified as the mean instantaneous firing rate during the hold phase.

### Skin clearing and immunostaining

Under deep anesthesia (Beuthanasia-D Special, 75 mg/kg, i.p.; Merck Animal Health, Kenilworth, NJ), a 3-mm full-thickness punch biopsy of footpad was collected from the plantar surface of the hindpaw. Tissue samples were immediately immersed in 4% paraformaldehyde (PFA) in 0.1 M phosphate-buffered saline (PBS, pH 7.4) and fixed overnight at 4°C with gentle agitation. Following fixation, samples were washed three times in PBS (10 minutes per wash) at room temperature to remove residual PFA prior to further processing.

Fixed samples were preserved using SHIELD reagents (LifeCanvas Technologies) per the manufacturer’s protocol and then cleared using a fDISCO+ protocol ^25^. All procedures were performed at 4°C with continuous shaking unless otherwise specified. Briefly, samples were washed sequentially through 50%, 70%, and 80% tetrahydrofuran (THF), then twice in 95% THF. Samples were delipidated in dichloromethane 100% DCM (Sigma-Aldrich), washed twice in 95% THF, then rehydrated through a descending THF series (80%, 70%, 50%). Residual solvent was removed by washing in 0.2% PBST followed by 1× PBS at room temperature. Fluorescence labeling with PGP9.5 (Proteintech Group Inc., IL; Cat# 17430) and secondary antibody in cleared samples were performed with eFLASH ^35^. For refractive index matching, samples were incubated sequentially in 50% EasyIndex (LifeCanvas, RI = 1.52) overnight at 37°C. Samples were finally transferred to EasyIndex at room temperature for refractive index matching and storing until imaging.

### 3D imaging and filament analysis

Three-dimensional images of cleared skin samples were acquired using a light-sheet fluorescence microscope (Alpenglow Bioscience and LifeCanvas Technologies). Z-stacks spanning the full tissue thickness were collected at a step size of 1.8μm for 3.6x magnification. Raw image stacks were imported into Imaris software (version 10.1.1; Oxford Instruments, Abingdon, UK) for visualization and quantitative analysis. Nerve fibers were semi-automatically reconstructed as three-dimensional filament objects using the Imaris Filament Tracer module, with tissue autofluorescence used to define the dermis-epidermis boundary via the surface module. The following parameters were extracted: (1) total length of free nerve ending per fiber, and (2) intraepidermal free nerve ending density (number of nerve endings per epidermal area). All filament reconstructions were reviewed and manually corrected by an investigator blinded to treatment group.

### Data analysis

Sample sizes for each experimental group were determined based on power analyses, relevant literature, and prior experience. All data are presented as mean ± standard error of the mean (SEM). Statistical analyses were conducted using GraphPad Prism 10 (GraphPad Software, La Jolla, CA). Data distributions were assessed for normality and homogeneity of variance; when these assumptions were not met, appropriate non-parametric tests were applied. Statistical comparisons among groups were performed using one-way ANOVA with Tukey’s post hoc tests, Chi-square test, or unpaired two-tailed t-tests, as appropriate. A significance threshold of P < 0.05 was applied to all tests. Significant differences are indicated in figures as: P < 0.05 (*), P < 0.01 (**), and P < 0.001 (***).

## RESULTS

### 1. Repeated paclitaxel treatment does not alter conduction velocity

Previous work reported that cisplatin reduces the overall conduction velocity (CV) of mouse tail nerves as measured from compound action potentials ^23^. To determine whether paclitaxel produces analogous effects at the single-fiber level, we examined CV in skin-nerve preparations harvested five weeks after paclitaxel treatment. Single-unit CV was calculated from the latency of electrically evoked action potentials and used to classify units as C-fibers (CV < 1.2 m/s) or Aδ-fibers (CV 1.2-10 m/s).

A greater proportion of fibers in the paclitaxel-treated group failed to generate an action potential in response to electrical stimulation compared to vehicle-treated animals, despite remaining responsive to mechanical stimuli applied to their receptive fields. In vehicle-treated animals, 8 of 380 recorded fibers (2.1%) were unresponsive to electrical stimulation, whereas in paclitaxel-treated animals, 31 of 340 fibers (9.1%) failed to respond electrically (**Fig. 1B**). Because mechanical stimuli activate fibers at greater tissue depth than electrical stimuli delivered via an electrode lowered into the superficial skin, this increase in electrically unresponsive fibers suggests that paclitaxel may impair axonal membrane excitability or disrupt structural integrity in a subset of peripheral nociceptors, consistent with the known axonopathic effects of taxane chemotherapeutics.

Among fibers successfully driven by electrical stimulation, CV values were plotted as scattergrams for each group (**Fig. 1C**). Consistent with prior reports ^34, 36^, repeated paclitaxel treatment did not significantly alter the mean CV of either C- or Aδ-fibers compared to vehicle-treated animals. The preservation of mean CV across groups confirms that the CV-based classification criteria remain valid for distinguishing fiber subtypes in both conditions, ensuring that subsequent between-group comparisons are not confounded by shifts in fiber classification.

A previous study suggested that paclitaxel induces PIPN by inhibiting Kv7 ion channels in primary sensory neurons ^33^. We therefore examined whether co-administration of the Kv7 channel activator retigabine affected CV. Retigabine was administered daily from Days 1 through 10, overlapping with the paclitaxel treatment window. The number of electrically non-responsive fibers was lower in the paclitaxel + retigabine group compared to the paclitaxel-alone group (**Fig. 1B**). Retigabine co-treatment did not alter the CV of C-fibers or Aδ-fibers relative to vehicle or paclitaxel-treated groups (**Fig. 1C**).

### 2. Repeated paclitaxel treatment increases nociceptor excitability

Prior sharp electrode recordings demonstrated that paclitaxel increases the proportion of non-nociceptors exhibiting spontaneous firing; however, this approach provides limited resolution for characterizing nociceptor-specific excitability changes ^36^. We therefore used skin-nerve preparations to directly assess whether paclitaxel enhances spontaneous activity in peripheral nociceptors.

In vehicle-treated animals, the majority of nociceptors were electrically silent at rest: spontaneous firing was observed in only 25.26% of C-fibers and 20.83% of Aδ-fibers (**Fig. 2A and B**). Paclitaxel treatment substantially shifted the excitability profile: 61.77% of C-fibers and 53.47% of Aδ-fibers from paclitaxel-treated animals exhibited spontaneous discharge, representing approximately 2.4- and 2.6-fold increases relative to vehicle-treated controls, respectively (**Fig. 2A and B**). These increases were statistically significant for both fiber types.

**Figure 2.**
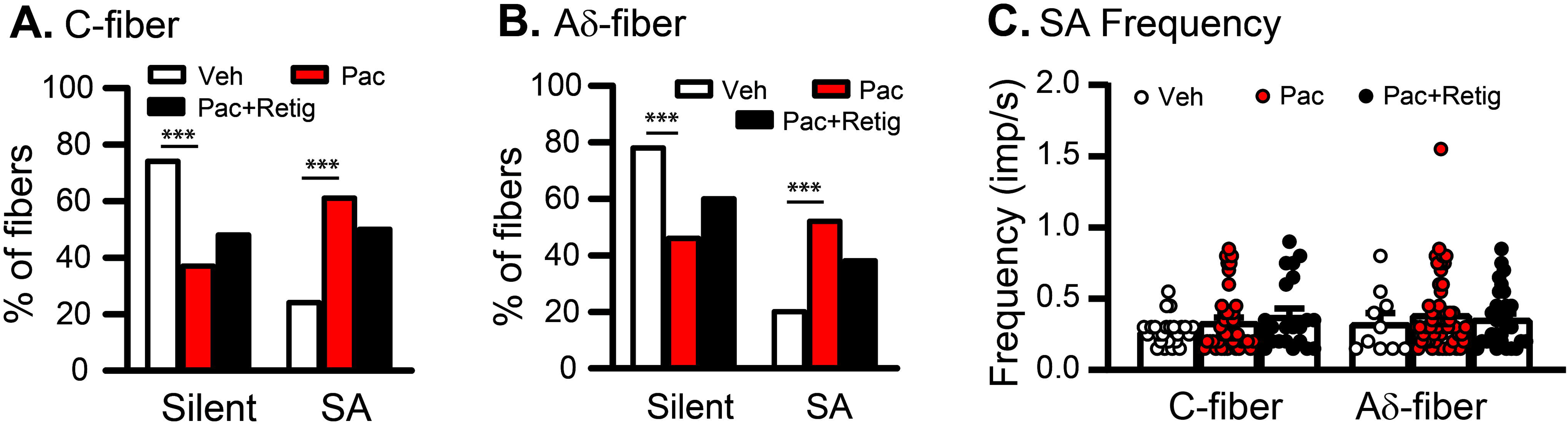
The effects of paclitaxel on the excitability of nociceptors. (**A**) Proportion of C-fibers from vehicle (n = 96), paclitaxel (n = 68), and paclitaxel+retigabine (n = 43) groups that are silent or displaying spontaneous firing. Chi-square test, X^2^ = 23.16, P < 0.0001. (**B**) Proportion of Aδ-fibers from vehicle (n = 42), paclitaxel (n = 131), and paclitaxel+retigabine (n = 136) groups that are silent or displaying spontaneous firing. Chi-square test, X^2^ = 14.65, P = 0.0007. (**C**) Frequency of Aδ- and C-fibers with spontaneous firing. Repeated measures one-way ANOVA followed by Tukey’s multiple comparison test (C-fibers, F_(2, 83)_ = 1.685, P = 0.1917; Veh vs Pac, P = 0.4340; Veh vs Pac+Retig, P = 0.1714. Aδ-fibers, F_(2, 93)_ = 0.39, P = 0.6782; Veh vs Pac, P = 0.7252, Veh vs Pac+Retig, P = 0.9358). Each dot in column represents one fiber. Veh, vehicle; Pac, paclitaxel; Retig, retigabine.

While paclitaxel increased the prevalence of spontaneous activity in both C- and Aδ-fibers, the frequency of spontaneous discharge among active fibers was not significantly altered in either fiber type compared to vehicle controls (**Fig. 2C**). These findings indicate that paclitaxel augments nociceptor excitability primarily by recruiting previously silent fibers into spontaneous activity, rather than by increasing the discharge rate of already-active fibers. This recruitment of peripheral nociceptor activity likely contributes to the spontaneous pain and mechanical hypersensitivity characteristic of PIPN.

Co-administration of retigabine reversed the paclitaxel-induced changes, reducing the proportion of spontaneously active fibers and restoring the proportion of silent fibers toward levels observed in vehicle-treated animals for both fiber types (**Fig. 2A and B**).

### 3. Repeated paclitaxel treatment decreases the mechanical threshold of C-fibers but not Aδ-fibers

We next characterized the mechanical response properties of individual nociceptors using a ramp-and-hold mechanical stimulation protocol. In skin-nerve preparations from vehicle-treated animals, Aδ- and C-fibers displayed comparable mechanical activation thresholds (112.21 ± 8.91 mN vs. 119.64 ± 5.45 mN; P = 0.4564). Paclitaxel treatment, however, differentially affected the two fiber types. Aδ-fiber mechanical thresholds remained unchanged relative to vehicle-treated animals, whereas C-fibers exhibited a significant reduction in mechanical threshold (119.64 ± 5.45 mN vs. 94.02 ± 7.24 mN; P = 0.0194; **Fig. 3B**), indicating selective peripheral sensitization of C-fibers to mechanical stimuli. Co-treatment with retigabine normalized C-fiber mechanical threshold (from 94.02 ± 7.24 mN to 120.16 ± 8.80 mN; P = 0.0492; **Fig. 3B**), further supporting a broad rescue of paclitaxel-induced nociceptor dysfunction by Kv7 channel activation.

**Figure 3.**
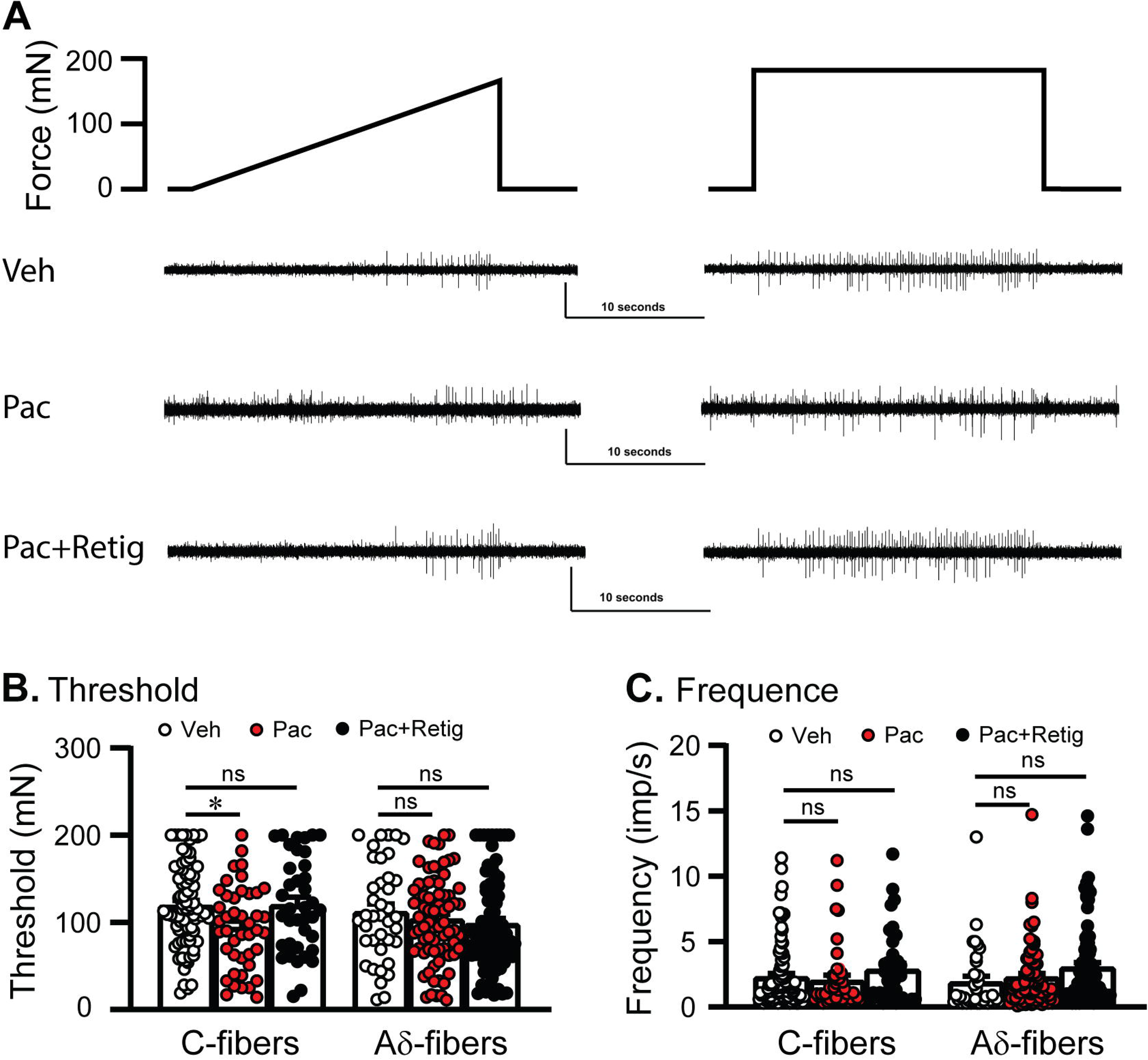
Sensitivity of nociceptors to mechanical stimulation. (**A**) Mechanical stimulation protocol and representative traces upon stimulation seen in C-fibers from Vehicle and Paclitaxel groups. The mechanical stimulation includes an ascending ramp phase (0-170 mN over 20 s) followed by a square wave pulse (180 mN, 10s). (**B**) The summary data comparing the mechanical thresholds of C- and Aδ-fibers between Vehicle and Pacclitaxel groups. C-fibers, F_(2, 156)_ = 4.262, P = 0.0157; Veh vs Pac, P = 0.0194; Veh vs Pac+Retig, P = 0.9985. Aδ-fibers, F_(2, 194)_ = 0.9054, P = 0.4046; Veh vs Pac, P = 0.6888, Veh vs Pac+Retig, P = 0.3738. (**C**) The effect of paclitaxel treatment on the discharge frequency of Aδ- and C-fibers upon suprathreshold stimulation. C-fibers, F(2, 171) = 1.231, P = 0.2946; Veh vs Pac, P = 0.8314; Veh vs Pac+Retig, P = 0.4496. Aδ-fibers, F(2, 191) = 2.655, P = 0.0729; Veh vs Pac, P = 0.7974, Veh vs Pac+Retig, P = 0.0.1033. Repeated measures one-way ANOVA followed by Tukey’s multiple comparison test. Veh, vehicle; Pac, paclitaxel; Retig, retigabine.

To further characterize the functional consequences of this sensitization, we delivered suprathreshold mechanical stimuli and quantified the evoked firing frequency. Paclitaxel did not significantly alter the suprathreshold firing frequency of Aδ-fibers (1.93 ± 0.42 imp/s vs. 2.29 ± 0.28 imp/s; P = 0.7974; **Fig. 3C**) or C-fibers (2.31 ± 0.26 imp/s vs. 2.04 ± 0.39 imp/s; P = 0.8314; **Fig. 3C**) compared to vehicle controls.

Together, these findings demonstrate that repeated paclitaxel treatment produces a selective reduction in C-fiber mechanical activation threshold while leaving Aδ-fiber mechanical responses largely intact. This fiber-type-specific sensitization provides a potential peripheral mechanism for the mechanical allodynia and hyperalgesia observed in PIPN.

### 4. Free nerve terminals are degenerated after paclitaxel treatment

In the rabbit corneal epithelium, C-fiber sensory terminals form vertically oriented processes running mostly perpendicular to the corneal surface, whereas Aδ-fiber terminals extend as horizontally oriented processes parallel to the epithelial surface ^18^. We therefore investigated whether C- and Aδ-fiber terminals show similar spatial organization in rat glabrous skin. Because conventional intraepidermal nerve fiber (IENF) analysis provides morphological information only for vertically oriented nerve endings at the dermal-epidermal junction, we employed fDISCO tissue clearing of glabrous skin to acquire three-dimensional images, enabling unbiased visualization of nerve ending architecture across treatment conditions.

In samples from all animals (n = 11), vertically oriented tree-like nerve fibers were readily observed, exhibiting continuous extension and branching that terminated as unencapsulated free nerve endings. Horizontally oriented fibers were observed in only one sample (**Supplementary data, Video S3**). In sham-treated animals, the majority of nerve fibers crossed the dermis-epidermis boundary into the epidermis. After paclitaxel treatment, however, terminal branch length was markedly decreased, with most free nerve endings terminating at or near the dermis-epidermis boundary rather than extending into the epidermis. Quantitative filament analysis revealed that the total terminal length per fiber was significantly reduced (**Fig. 4A** and **B**), along with a pronounced decrease in free nerve ending density in the epidermis (**Fig. 4C**). Co-treatment with retigabine substantially preserved fiber continuity and reduced the extent of terminal disruption compared to the paclitaxel-alone group.

**Figure 4.**
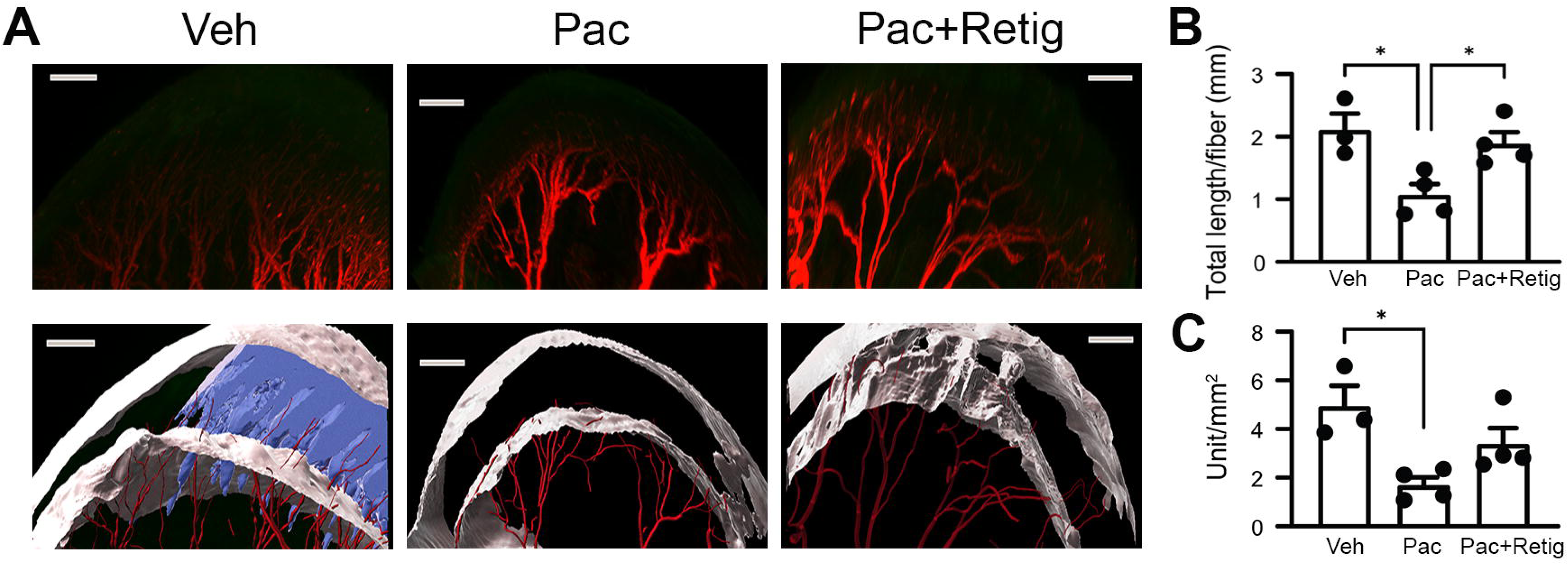
The effects of treatments on the integrity of nerve endings in glabrous skin. (**A**) Representative images showing representative 3D images and Imaris filament analysis of vehicle-, paclitaxel-, and paclitaxel+retigabine-treated samples. Scale bar 300 μm. (B) Summary data indicating the total length of free nerve ending per fiber. F(2,8) = 7.667, P = 0.0138, Veh vs Pac, P = 0.0175; Veh vs Pac+Retig, P = 0.7396; Pac vs Pac+Retig, P = 0.0373. One-way ANOVA. Dots in each column represent an animal. (C) Summary data indicates the quantification of the density of free nerve endings in epidermis. F(2,8) = 7.072, P = 0.017, Veh vs Pac, P = 0.0141; Veh vs Pac+Retig, P = 0.2361; Pac vs Pac+Retig, P = 0.1486. One-way ANOVA. Dots in each column represent an animal. Dots in each column represent an animal. *, P < 0.05; Veh, vehicle; Pac, paclitaxel; Retig, retigabine.

## DISCUSSION

Chemotherapy-induced peripheral neuropathy presents with characteristic “stocking-and-glove” sensory deficits, and morphological studies have documented pronounced degeneration of peripheral sensory terminals ^15, 30, 33^. Yet many electrophysiological investigations of chemotherapy-induced sensory dysfunction have focused on DRG somata or sural nerves, sites where overt morphological changes are typically absent ^34, 36^. Moreover, *ex vivo* skin-nerve recordings have suggested that the overall number of C-fibers remains largely unchanged following chemotherapy ^29^. In contrast, our combined electrophysiological and 3D morphological analyses provide converging evidence that both C- and Aδ-fibers in the peripheral skin undergo functional alterations following paclitaxel treatment, accompanied by significant degeneration of C-fiber terminals at the epidermal junction. These findings support a direct role for terminal pathology in chemotherapy-induced peripheral neuropathy.

Many Aβ-fibers have CVs in the Aδ range, but their mean CV is significantly slower than that of Aδ-fibers, and these slower Aβ-fibers are typically low-threshold mechanoreceptors ^9^. In our study, the CVs of Aδ-fibers from all three groups clustered around the slower peak, suggesting that these fibers were nociceptors. All recorded Aδ-fibers also responded to noxious mechanical stimulation, consistent with nociceptor identity. It is possible that some Aδ-fibers were sensitized by paclitaxel treatment to respond to non-noxious stimuli. Such sensitized nociceptors could have been excluded from further recording under our selection criteria, potentially leading to underestimation of paclitaxel’s effects on Aδ-fiber mechanical thresholds.

Previous DRG soma and sural nerve recordings at the popliteal fossa also showed that paclitaxel treatment increases the number of nociceptors with spontaneous discharge ^34, 36^. Our skin-nerve recordings demonstrate that spontaneous discharge after paclitaxel originates from nerves in the hindpaw itself, possibly arising from intact axons whose sensory terminal arbors have been partly or completely damaged. Spontaneous discharge can be recorded in intact C-fibers traveling alongside degenerated myelinated axons in rats and monkeys ^3, 32^, but this activity has a distinctly low frequency (approximately one impulse per minute), well below the firing rates of the spontaneously active fibers in our study and in sural nerve recordings (at least 6-fold faster). The chemotherapy-evoked C- and Aδ-fiber discharge therefore most likely originates within or near the fibers’ sensory terminal arbors, though we cannot exclude a contribution from the length of axon between the terminals and the recording site.

Spontaneous discharges in compressed DRG neurons are classified into Type I, which depends on subthreshold membrane potential oscillations (SMPOs) and originates in the soma, and Type II, which is oscillation-independent and likely originates in the axon ^17^. In DRG somata, action potentials consistently emerge from the depolarizing phase of oscillations, indicating a causal relationship between oscillations and spiking ^8, 24^. Chronic paclitaxel treatment depolarizes DRG somata, promoting and amplifying SMPOs and thereby increasing the incidence and frequency of spontaneous discharge. Type II discharge, by contrast, is membrane potential-independent ^17^. In our skin-nerve recordings, we observed increased incidence of axonal ectopic discharge in nociceptors without a corresponding change in discharge rate. The terminal structural disruptions revealed by 3D imaging may directly create ectopic axonal pacemaker sites, providing a substrate for Type II-like spontaneous discharge. Although the two types are distinct, spikes generated at multiple sources can interact to produce complex discharge patterns ^4, 17^, and axonally generated Type II activity can serve as an additional depolarizing input to the soma, amplifying the overall ectopic barrage under pathological conditions.

Paclitaxel directly excites DRG neurons, including nociceptors, through Kv7 ion channels ^33^, but the extent to which this mechanism contributes to nociceptor hyperexcitability at the time point of our recordings (five weeks post-treatment) is uncertain, since the level of paclitaxel remaining in axons at this stage is unknown. Piezo2, a rapidly adapting mechanically activated ion channel, is responsible for mechanosensitivity in most low-threshold mechanoreceptor subtypes involved in innocuous touch ^26^. However, recent work has revised this picture: optogenetic activation of Piezo2-expressing sensory neurons induces nociception in mice, and skin-nerve recordings from Piezo2 knockout mice show diminished Aδ-nociceptor and C-fiber firing in response to mechanical stimulation ^22^. Alternatively, Piezo1 in keratinocytes has been implicated in paclitaxel-induced mechanical sensitization ^21^. In our study, however, only C-fibers showed a decreased mechanical threshold after paclitaxel. TRPV4 is also mechanosensitive ^11, 19^, expressed in nociceptors ^28^, and mediates paclitaxel-induced mechanical hyperalgesia through PKC-epsilon activation ^2, 7^. Notably, electrophysiological recordings from primary bladder afferents and knee joint afferents show that only C-fibers, not Aδ-fibers, are influenced by TRPV4 antagonists during both normal and inflammatory pain ^1, 27^. The selective C-fiber mechanical threshold reduction we observed suggests that paclitaxel-induced sensitization of TRPV4 may be the primary contributor to mechanical sensitization in PIPN.

Nerve fibers in rat glabrous skin form vertically oriented, tree-like structures that branch continuously and terminate as unencapsulated free nerve endings. This contrasts sharply with the rabbit corneal epithelium ^18^, where Aδ-fibers extend horizontally whereas C fibers form clustered vertical terminals. The architectural difference is probably linked to their functions. In the cornea, the horizontal Aδ-fiber layout functions as a planar detection grid optimized for absolute protection. The cornea does not need to discriminate gradations of force. Sub-millinewton stimuli uniformly triggers intense pain and protective blinking, and the architecture reflects that priority. Glabrous skin has a different problem to solve. It must handle constant, varied mechanical loads during active exploration and distinguish a gentle touch from a noxious puncture. The vertical, tree-like organization is well suited to this task. A single parent axon samples multiple tissue depths simultaneously, and as increasing force deforms progressively deeper strata, terminal recruitment scales accordingly. This arrangement gives the CNS a way to estimate mechanical intensity from population-level activity along a vertical axis. The overlapping vertical networks also retain enough redundancy to preserve high-threshold mechanical sensation, and may even become sensitized when mechanotransducers within this network are upregulated, even as C-fiber free nerve ending density falls after chemotherapy.

Several limitations should be noted. First, while 3D imaging revealed the full architecture of sensory nerve fibers in glabrous skin and their treatment-related changes, we did not distinguish A-fibers from C-fibers morphologically. Future work incorporating myelin markers (such as Myelin Protein Zero) will be needed to determine whether these two populations show distinct structural responses to chemotherapy. Second, we restricted the study to male rats to eliminate variability from estrous cycle stage. This is a reasonable simplification because paclitaxel-induced mechanical allodynia develops similarly in both sexes ^14^. The alterations observed here are therefore likely to generalize to females, though direct testing remains warranted.

## Supporting information

Supplemental Data Video S1 (sham)

Supplemental Data Video S2 (Paclitaxel)

Supplemental Data Video S3 (Paclitaxel+retigabine0)

## Footnotes

The project was supported by grants from National Institutes of Health (R01 CA 273001, CA208765 to Q.Y., and DC016328 to Z.W.), Craig H. Neilsen Foundation (383428 to Q.Y.), and Department of Defense Hearing Restoration Research Program (RH220042 to Z.W.). The authors declare no competing financial interests.

## List of Nonstandard Abbreviations

PIPN: Paclitaxel-induced peripheral neuropathy
DRG: Dorsal root ganglia
IENFs: Intraepidermal nerve fibers
CV: Conduction velocity
fDISCO: Immunolabeling-enabled three-dimensional imaging of solvent-cleared organs
DCM: Dichloromethane
PGP9.5: Protein gene product 9.5
SMPOs: Subthreshold membrane potential oscillations

## SUPPLEMENTARY DATA

**Video S1:** Light-sheet 3D image of cleared footpad from vehicle-treated rat. Tissue was labeled with PGP9.5 (red) and reconstructed with the Imaris filament module. Related to Figure 4.

**Video S2:** Light-sheet 3D image of cleared footpad from paclitaxel-treated rat. Tissue was labeled with PGP9.5 (red) and reconstructed with the Imaris filament module. Related to Figure 4.

**Video S3:** Light-sheet 3D image of cleared footpad from paclitaxel + retigabine co-treated rat. Tissue was labeled with PGP9.5 (red) and reconstructed with the Imaris filament module. Related to Figure 4.

